# Use of botanical gardens as arks for conserving pollinators and plant-pollinator interactions: A case study from the US Northern Great Plains

**DOI:** 10.1101/2021.02.26.433109

**Authors:** Isabela B. Vilella-Arnizaut, Charles B. Fenster

## Abstract

Botanical gardens have contributed to plant conservation through the maintenance of both living and preserved plant specimens for decades. However, there is still a large gap in the literature with regards to understanding the potential conservation value botanical gardens could provide for local pollinators. We investigated how plant-pollinator community structure and diversity may differ between botanical gardens and native habitats by sampling and comparing between two environments: a restored native grassland patch within a local botanical garden and fifteen native, remnant temperate grassland sites in the Northern Great Plains. We found pollinator diversity within the restored native grassland patch was greater than 55% of total remnant temperate grassland transects throughout the entire flowering season, while plant diversity and network community metrics between the two environments remained similar throughout, except that remnant prairies have more links (higher connectance) with pollinators than the garden patch. Overall, our findings demonstrate the promising role restored native grassland patches in botanical gardens could play as reservoirs for local pollinator communities by supporting plant-pollinator interactions comparable to those found in native habitat remnants in the same region.

## Introduction

Biodiversity loss is a global crisis that many countries have attempted to address through numerous methods of preservation and conservation management strategies (Mutia 2009; Benedict and McMahon 2006; Hostetler et al. 2011; Bortree et al. 2013). The propagation and maintenance of botanical gardens is one strategy that has been implemented, particularly for plant conservation (Hurka 1994; Primack and Miller-Rushing 2008; Powledge 2011; Miller et al. 2016; Chen and Sun 2018). Botanical gardens and urban green spaces may serve as potential reservoirs for pollinators (Pinheiro et al. 2006; Levé et al. 2019; Buchholz et al. 2020). However, there is still a gap in the literature with regards to understanding how botanical gardens support pollinators and preserve plant-pollinator interactions. For example, a literature search (Web of Science, November 11^th^, 2020) using the terms “botanical gardens and pollinator diversity” and “botanical gardens and plant diversity” resulted in 14 and 293 citations, respectively, demonstrated much greater focus on the contribution of botanical gardens to plant conservation and diversity than pollinators. Clearly, the potential conservation value botanical gardens hold could extend beyond plant conservation. These gardens could provide space for several resources that pollinators utilize (i.e., foraging and nesting resources), even in areas that would typically be considered resource-poor (e.g., cities) (Lewis et al. 2019).

With approximately 1,775 botanical gardens worldwide (Botanic Gardens Conservation International, 2020), these sites could provide increasingly important conservation resources that can be utilized to alleviate the accumulating threats towards pollinators (i.e., habitat loss and fragmentation, pesticide use, and invasive species introductions) (Kearns et al. 1998; Kremen et al. 2002; Steffan-Dewenter et al. 2005). Habitat loss and fragmentation are two primary causes for pollinator decline (Potts et al. 2010; Vanbergen et al. 2013; Habel et al. 2019) and are expected to continue with increased urbanization and agricultural intensification (Foley et al. 2005; Lundgren and Fausti 2015; UN DESA, 2018). This is particularly concerning considering animal-driven pollination is essential to the reproduction of over 70% of flowering plant species (Potts et al. 2010) and 35% of crops globally (Klein et al. 2007; Vanbergen et al. 2013).

With the space and habitat that is left, can we look to botanical gardens as a proxy for native habitat to provide refugia for pollinators? Urban green spaces and botanical gardens can positively influence pollinator abundance or diversity depending on total area, floral abundance, and degree of urbanization (Tommasi et al. 2004; Gotlieb et al. 2011; Fortel et al. 2014; Micholap et al. 2017). In the United States, there are even cities which support a greater diversity of native bees than neighboring rural areas (Hall et al. 2017; US Fish and Wildlife Service, 2015). Furthermore, there is a rise in initiatives to promote expanding urban private and public garden space with the hopes of promoting and sustaining stable pollinator communities (e.g., The Million Pollinator Garden Challenge sponsored in part by the United States Botanic Garden Conservatory). With the increased interest in carving out urban spaces for pollinators, there is a need to assess the stability of plant-pollinator community structures in the context of botanical gardens (Spiesman and Inouye 2013). The stability of pollination services is dependent upon maintaining diverse and resilient plant-pollinator communities (Klein et al. 2007). Network theory has been utilized to examine how the mutualistic interactions within plant-pollinator communities influences their structure and in a broader sense, interpret the mechanisms behind biodiversity and community resilience (Memmott et al. 2004; Bascompte et al. 2006; Blüthgen et al. 2008; Dupont et al. 2009; Hadley and Betts 2012; Spiesman and Inouye 2013; Soares et al. 2017; Redhead et al. 2018). Using a network-based approach, we can assess how plant-pollinator communities are structured in botanical gardens to determine if they may serve as supplementary resources for preserving plant-pollinator interactions. However, we lack information on how plant-pollinator interactions in botanical gardens compare to nearby natural habitat.

We focus our study in McCrory Gardens, a botanical garden located in Brookings, (eastern) South Dakota, a small city with a population of 24,000. Our goal is to assess the structure of plant-pollinator communities in botanical gardens and how they compare to nearby natural habitats. The garden is located within the Prairie Coteau, a region containing some of the largest remaining tracts of tall-grass habitat in the Northern Great Plains, with temperate grassland remnants nestled within an actively transforming and working landscape (Bauman et al. 2016). In the center of McCrory Gardens, we focused our sampling within a 1,600 m^2^ area designated as a restored native grassland patch that was established in 2018. This planted native grassland garden, embedded within a larger landscape of varying patches of natural and modified habitat, provides us a study system with which to compare how plant-pollinator communities within botanical gardens measure to those found in natural remnant habitats. Habitat loss and fragmentation is still a substantial threat to the temperate grasslands of the Northern Great Plains with documented rates of conversion from grassland to agricultural crops reaching ∼1.0-5.4% annually from 2006 to 2011 (Wright and Wimberly 2013). A better understanding of plant-pollinator community structure in botanical gardens and their role in pollinator conservation will become increasingly important for future management decisions seeking to bolster pollination services.

We measure the diversity of plant-pollinator communities within natural temperate grassland areas and a restored grassland patch in a botanical garden, then quantify plant-pollinator interactions using a network-based approach in order to answer the following questions: 1) How does the pollinator diversity found within a restored grassland patch located in a botanical garden compare to the diversity found within native temperate grassland sites? 2) Likewise, how does the diversity of the biotically-pollinated plant community within a restored grassland patch compare to that of native temperate grassland sites? And 3) What is the overall structure of plant-pollinator community interactions within a restored grassland patch located in a botanical garden and how do they compare on average to plant-pollinator networks in native temperate grassland sites? These questions become increasingly relevant with the progressive loss of biodiversity as urbanization and agricultural intensification continues to encroach upon natural landscapes (Ramankutty et al. 1999; Hoekstra et al. 2005).

## Materials and Methods

### Study area

McCrory Gardens (long: −96.791080, lat: 44.309100) is a botanical garden located in Brookings, South Dakota, that is operated and maintained by South Dakota State University (SDSU). The garden is located 300 m from an 18-hectare SDSU agricultural plot to the north and about 2 km from private farmland to the east. Otherwise, it is surrounded by the SDSU campus, residential housing and apartments, shopping malls, large box stores, and major thoroughfares. Founded in the early 1960s, McCrory Gardens contains over 10 hectares of display gardens that showcases a variety of ornamental and native plant species. The garden’s origin began with a mission to maintain a research garden that displays and educates the public on plant species that were or are a part of the South Dakota landscape. In continuation with this original mission statement, the Prairie Centennial Garden was established in 2018 in the center of McCrory Gardens. This 1,600 m^2^ plot is a restoration native grassland garden with 85% of the plants grown from seed by the McCrory gardens staff (seeds provided by Prairie Moon Nursery in Winona, MN & Jelitto Perennial Seeds) and the remaining 15% of plants were relocated or reused from other areas within McCrory Gardens. Seed from Jelitto was not locally sourced but came from locations as close as Minnesota and as far as Colorado.

To compare the diversity of insect pollinators and plants, and plant-pollinator community structure between the botanical garden and native temperate grassland remnants, we selected fifteen remnant temperate grassland sites within the Prairie Coteau region in eastern South Dakota. Within South Dakota, this region covers approximately 17 counties and contains some of the most land cover remaining of native tall-grass prairie in the Northern Great Plains (Bauman et al. 2016). In eastern South Dakota, approximately 17% of the undisturbed grasslands within the Prairie Coteau region remain intact making this a valuable resource for tall-grass habitat in the Northern Great Plains. Remnant temperate grassland sites ranged in size from 8 to > 400 hectares and were selected based on quality of the site as advised by local experts and managers (see *Acknowledgements*), as well as manifesting a range of site characteristics, including size, local landscape use, and proximity to other semi-disturbed grasslands. Full description of site names, location coordinates, county, size and ownership are provided in Vilella-Arnizaut et al. (bioRxiv, doi.org/10.1101/2021.02.12.431025).

### Data collection

#### Pollinator observations

We conducted pollinator observations in the restored native grassland patch in McCrory Gardens and fifteen remnant temperate grassland sites within eastern South Dakota between May and October 2019. We sampled a total of 10 transects within the restored native grassland patch in McCrory Gardens and 114 transects across all remnant temperate grassland sites throughout the entire growing season for one year, see Vilella-Arnizaut et al. (bioRxiv, doi.org/10.1101/2021.02.12.431025) for more details.

Pollinator observations were conducted for 30 minutes along 30 x 1 m transects on days warm enough to allow insect flight and in time periods when pollinators are expected to be active (15-35⁰ C, between 08:00 - 17:00 hours). We divided the sampling into three seasons: early (May-June), mid (July-August) and late (September-October). Season intervals were selected based on consistent flowering phenology shifts found in the plant communities of the Prairie Coteau. For example, species belonging to the genera *Anemone* (*Ranunculaceae*), *Viola (Violaceae)*, and *Sisyrinchium (Iridaceae)* predominately bloomed in the early season, while the mid and late seasons were dominated by species in the *Fabaceae* and *Asteraceae* (legume and sunflower families, respectively). Though these two families were predominately found in both seasons, the mid-season is distinct as this period marked a peak in the number of families in bloom with approximately six times more families present in our surveys in comparison to other seasons. Late season was characterized by a distinct shift in floral composition in which Asteraceae became the most prominent family in all sites with nearly all other families no longer flowering for the year.

From our roster of sites, which included the restored native grassland patch in McCrory Gardens and the fifteen temperate grassland sites, we randomly sampled each site until flowering ceased at each location. Location and direction of transects were randomized at each visit using a list of randomly generated numbers to determine number of steps and cardinal directions before placing transects down. Transects were geospatial referenced using a Trimble Geo 7x GPS unit with 1-100 cm accuracy. We walked the entire length of the transect and recorded all plant-pollinator interactions from within one meter of the transect line on both sides. We defined pollinators as insect floral visitors that made contact with both the male and female reproductive parts of the flower, a commonly used criterion (Fenster et al. 2004). We documented each pollinator and the associated biotically-pollinated plant species when an interaction occurred. Additionally, we documented pollinator return visits to plants.

Pollinators were identified in situ to family and genus, then to morphospecies in order to quantify insect diversity. The pollinator observations in our study only focus on diurnal pollinators, however, this does not present a significant bias in our sampling. Our data set portrays a robust, representative sample of the plant-pollinator networks in this region considering only one species (*Silene vulgaris*) detected in our floral surveys (described in the next section) relies on nocturnal pollination, and this one species was only present in 1 transect of the 124 sampled. Insect voucher specimens were collected in the field with an aspirator and net, later identified to lowest taxonomic level and then categorized into functional groups. Specimens were identified using resources available through discoverlife.org, bugguide.net, and Key to the Genera of Nearctic Syrphidae (Miranda et al. 2013). Voucher insect specimens were verified for sampling completeness using the help of experts and the Severin-McDaniel Insect Research Collection available at South Dakota State University.

### Floral surveys

Floral surveys were conducted directly after insect pollinator observation surveys along the same transect with a 1 m^2^ quadrat. The quadrat was placed at each meter mark from 0 to 30 m where we documented the presence of each biotically-pollinated plant species, number of individuals per species, number of flowering units defined as a unit of one (e.g., Ranunculaceae) or a blossom (e.g., Asteraceae) requiring a small pollinator to fly in order to access another flowering unit per species, and percent cover within quadrat per species. We also quantified the symmetry of flowers (radial vs. bilateral), since symmetry is often related to the degree of pollinator specialization (Fenster et al. 2004, Fenster and Marten-Rodriguez 2007). Hence, a greater proportion of either may affect the parameters of network analyses at the remnant temperate grassland communities and the restored native grassland patch in McCrory Gardens. Plant voucher specimens were not collected in McCrory Gardens, but photographs were taken and then verified by head gardener, Chris Schlenker. In the remnant temperate grassland sites, plant voucher specimens were collected and identified using Van Bruggen (1985) and verified with the help of experts (see *Acknowledgements*) and are curated at the C. A. Taylor Herbarium at South Dakota State University. Digitized plant collections for this study may be accessed on the Consortium of Northern Great Plains Herbaria (https://ngpherbaria.org/portal/).

### Pollinator and plant diversity

Pollinator and plant Shannon diversity were calculated using the ‘vegan’ package, version 3.6.3, in R (R Core Team 2013; Oksanen et al. 2019). The Shannon index is calculated using the following formula:

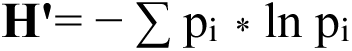

Where p_i_ is the proportions of each species found in a community and ln is the natural log. All diversity indices were natural log transformed. Shannon is the only diversity index presented in this study as it accounts for abundance, richness, and evenness. Pollinator diversity was calculated at the functional, family, and genus level by transect and season. Values used in pollinator diversity did not include return visits recorded during observation surveys. Although we recorded morphospecies in the field, we found genus to be the lowest, most robust taxonomic level in the data set for insect pollinators that could be identified with accuracy. Approximately 99.5% of total insect pollinator samples collected and observed within McCrory Gardens were identified to genus. Samples from McCrory Gardens that were not identified to genus were identified to next taxonomic level (family or functional group 0.5%) and given a catch-all classification which was used for pollinator genus analyses. Insect pollinators that could not be identified to genus were placed in a catch-all genus that consisted of the first five letters of their family name. For example, for a fly pollinator in the family Muscidae, we created a genus named Gen_Musci in the dataset in order to include these visitors in the analyses. We found pollinator genus diversity to be correlated with functional group diversity (Fenster et al. 2004) and family diversity in both environments (Supplemental table 1). We focus our results and comparisons on pollinator genus diversity to reduce the number of analyses. However, we provide the distribution data for all three categories of pollinator diversity for completeness. Statistical analyses for the remaining pollinator categories across seasons are provided in supplementary materials (Table 2).

Likewise, plant diversity was calculated at the family, genus, and species level by transect and season using number of individuals recorded during floral surveys. We generated correlation plots for all plant diversity levels as well and found that plant species diversity was correlated with family diversity and genus diversity (Supplemental tables 1). Thus, we focus on plant species diversity in our results and comparisons, but as above, we provide distribution data for the three categories of plant diversity for completeness. Statistical analyses for the remaining plant categories across seasons are provided in supplementary materials (Table 2). Details for correlation results and diversity metrics for both plant and pollinator communities in the natural temperate grassland sites are provided in Vilella-Arnizaut et al. (bioRxiv, doi.org/10.1101/2021.02.12.431025). However, we provide a synopsis of the diversity for the plant and pollinator communities in the natural temperate grassland sites in the results.

### Network analysis

We built quantitative visitation networks for each site using transects as our replicates to quantify plant-pollinator community structure. We calculated network metrics for each transect in order to statistically compare between the restored native grassland patch in McCrory Gardens and all remnant temperate grassland communities. We used transects as our replicates to compare network metrics between the two environments. We construct our network analysis based on the entire flowering season (May – October) because of limited sampling in the early and late seasons in both remnant temperate grassland sites and the garden. Networks were constructed using a matrix of interactions between plants and pollinators including unique and return visits recorded during pollinator observation surveys. Documenting return visits allows us to quantify plant-pollinator communities using weighted network values that also account for visitation frequency. For each network, we calculated network specialization (H2’), connectance and nestedness. We also provide the means of each network metric within a given season using transects as our replicates for the restored native grassland patch within McCrory Gardens and all remnant temperate grassland sites in supplementary materials (Table 3). All network metrics were calculated using the ‘bipartite’ package in R (Dormann et al. 2009).

### Statistical Analyses

We implemented Mann-Whitney U tests to compare pollinator genus diversity, plant species diversity and network metrics between remnant temperate grassland sites and the restored native grassland patch in McCrory Gardens using transects as our replicates. We compared these metrics across all three seasons since early and late season sampling were limited due to inclement flooding conditions. We used the ‘stats’ package in R to execute the Mann-Whitney U tests (R Core Team 2020). Chi-square tests were implemented in Microsoft Excel to examine differences in floral morphology (radial vs. bilateral symmetry) between the restored native grassland patch in McCrory Gardens and remnant temperate grassland communities.

## Results

### Pollinator community

Within the restored native grassland patch in McCrory Gardens, we observed 10 functional groups, 25 families, and 48 genera of pollinating insects (see Appendix A for full list). Among all 15 remnant temperate grassland communities, we observed 10 functional groups, 45 families and 79 genera of pollinating insects (see Vilella-Arnizaut et al. bioRxiv, doi.org/10.1101/2021.02.12.431025 for full list). Across all three seasons, we found a significant difference in pollinator genus diversity between environments with the restored native grassland patch in McCrory Gardens manifesting higher pollinator diversity (Genus: U =993, n1 = 114, n2 =10, p < 0.0002, Mean ± SE: Grassland Remnants =1.19 ± 0.05, Garden =1.85 ± 0.10). Shannon diversity of pollinator genera within the restored native grassland patch ranged from 1.28 - 2.28, while the Shannon diversity of pollinator genera within remnant temperate grassland sites ranged from 0 – 2.31 across all three seasons (Fig. 1). Early season pollinator genus diversity within the restored native grassland patch was 1.77 with Syrphidae (54%), Muscidae (15%) and Vespidae (12%) comprising the majority of observations (Fig. 2). Within remnant temperate grassland sites, early season pollinator genus diversity ranged from 0 – 2.03 with Syrphidae (36%), Muscidae (20%), Chloropidae (14.5%), and Halictidae (14%) dominating most interactions. Mid-season pollinator genus diversity within the restored native grassland patch ranged from 1.28 - 2.28 with Syrphidae (31%), Cantharidae (20%) and Tachinidae (9%) comprising the most observed interactions (Fig. 2). Mid-season pollinator genus diversity within remnant temperate grassland sites ranged from 0 – 2.31 with Apidae (27%), Syrphidae (25.7%), Cantharidae (12.5%), and Halictidae (11%) as the most common pollinators. During the late season, pollinator genus diversity within the restored native grassland patch in McCrory Gardens was 2.10 with Apidae and Syrphidae constituting nearly all interactions during this season with 68 and 25 percent, respectively (Fig. 2). Within remnant temperate grassland sites, late season pollinator genus diversity ranged from 0.3 – 1.9 with Syrphidae and Halictidae constituting 58 and 28 percent of observations, respectively.

**Figure 1.**
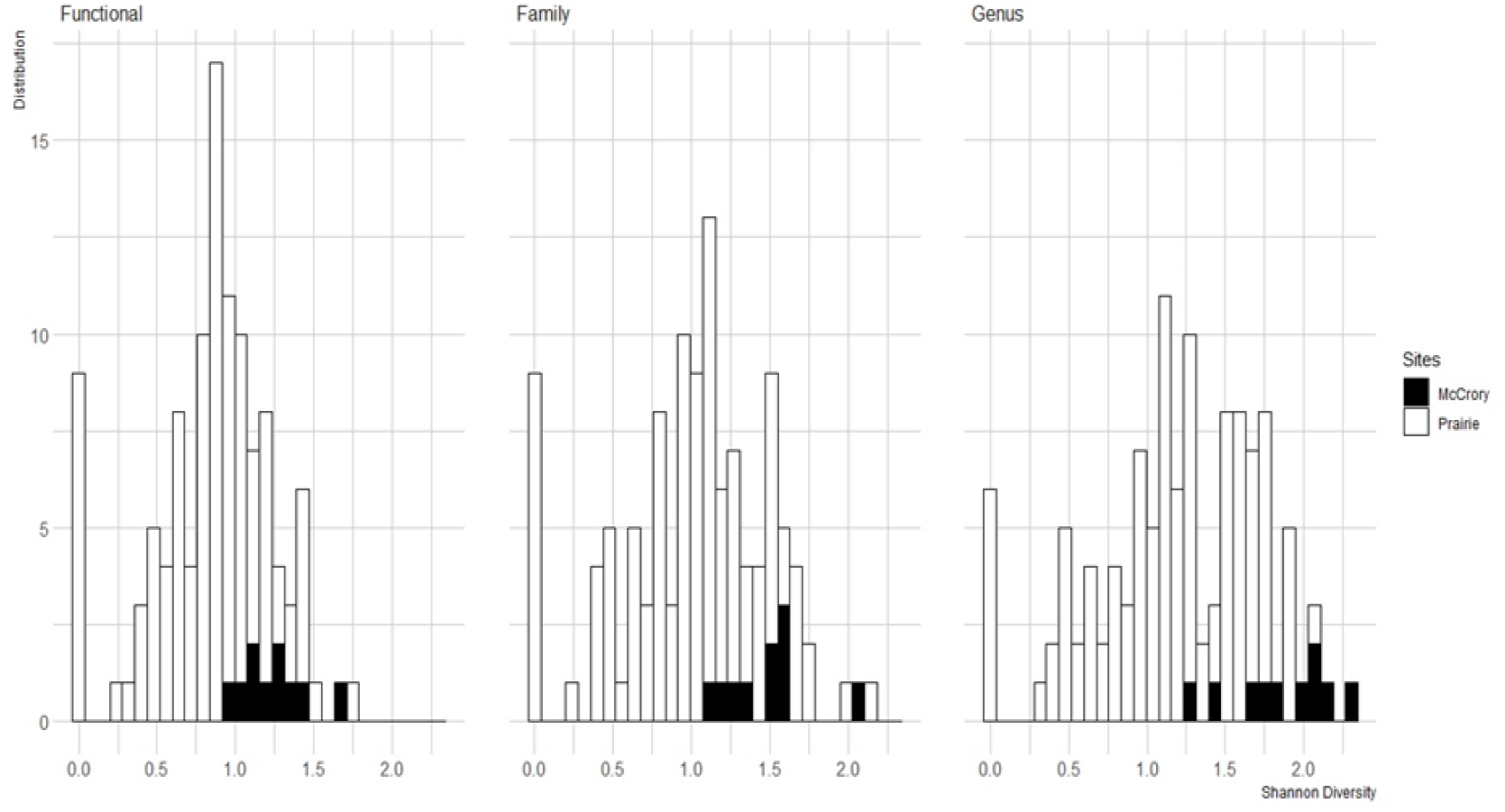
Distribution of Shannon diversity of pollinators at the functional, familial, and genera level between remnant temperate grassland sites in the Prairie Coteau near Brookings, SD and McCrory Gardens across the three sampling seasons (May – October 2019). The black bars refer to transects conducted in McCrory Gardens while the white bars refer to transects conducted in the remnant temperate grassland sites. Distributions demonstrate diversity on the transect level and are overlayed (i.e., not stacked) for comparison between sites. Height of distribution bars refer to number of samples (transects) that fell within diversity range indicated on the x-axis. Diversity displayed for remnant temperate grassland sites are also provided in Vilella-Arnizaut et al. (bioRxiv, doi.org/10.1101/2021.02.12.431025).

**Figure 2.**
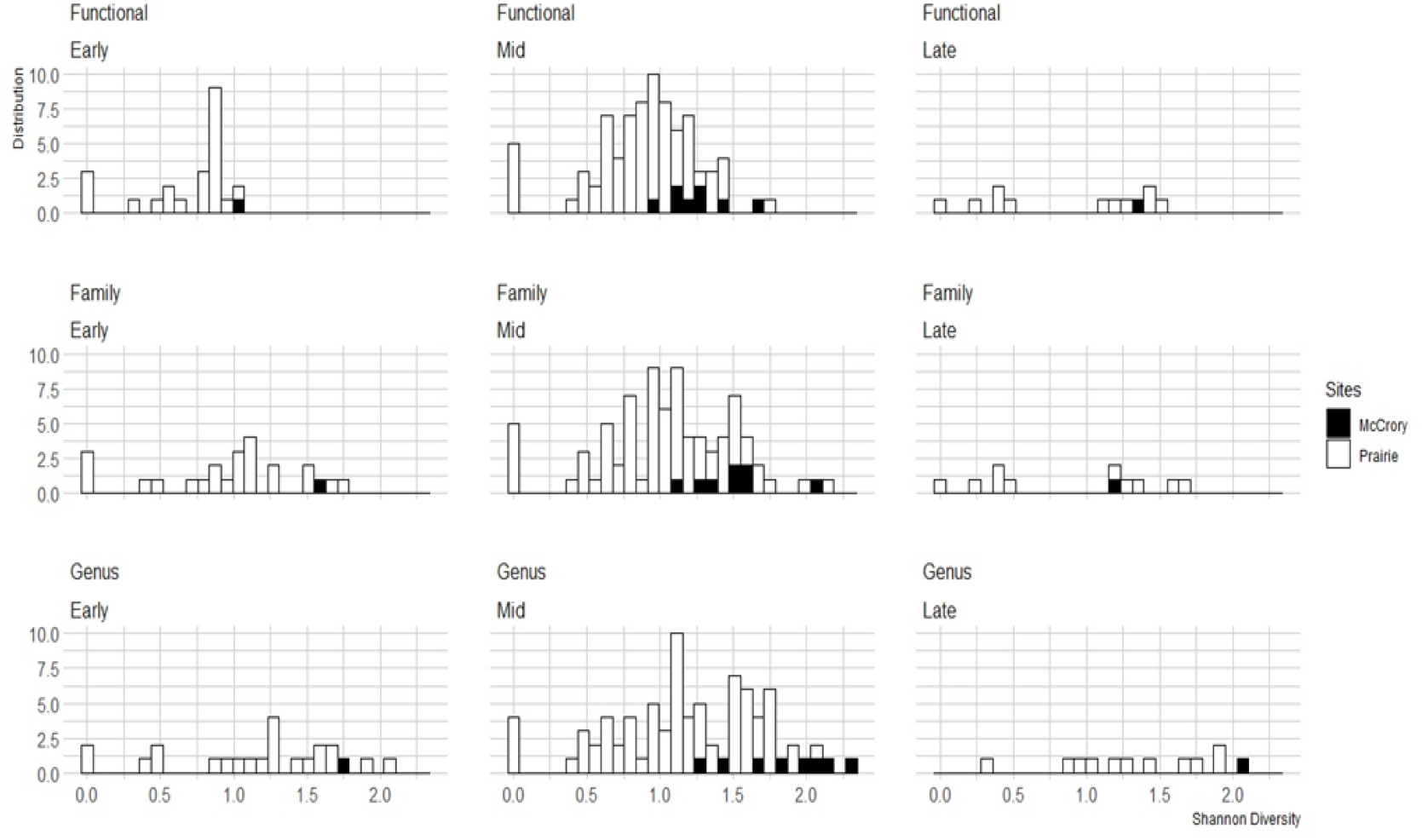
Comparing distribution of Shannon diversity of pollinators at the functional, familial, and genera level between remnant temperate grassland sites in the Prairie Coteau near Brookings, SD and McCrory Gardens for each sampling season in 2019 (Early: May – June, Mid: July – August, Late: September – October). The black bars refer to transects conducted in McCrory Gardens while the white bars refer to transects conducted in the remnant temperate grassland sites. Distributions demonstrate diversity on the transect level and are overlayed (i.e., not stacked) for comparison between sites. Height of distribution bars refer to number of samples (transects) that fell within diversity range indicated on the x-axis. Diversity displayed for remnant temperate grassland sites are also provided in Vilella-Arnizaut et al. (bioRxiv, doi.org/10.1101/2021.02.12.431025).

### Plant community

We sampled a total of 7 families, 19 genera, and 23 species of biotically-pollinated plants within the restored native grassland patch in McCrory Gardens (see Appendix B for full list). Among all 15 remnant temperate grassland communities, we sampled a total of 24 families, 61 genera, and 87 plant species (see Vilella-Arnizaut et al., bioRxiv, doi.org/10.1101/2021.02.12.431025 for full list). Out of 23 biotically-pollinated plant species in the restored native grassland patch, we determined 4 species displayed bilateral symmetry while 19 species displayed radial symmetry. Likewise, across remnant temperate grassland communities, we determined 25 species exhibited bilateral symmetry while 62 species exhibited radial symmetry. After conducting a chi-square test, we found no difference between environments with regards to proportion of floral morphology, χ^2^ (1 df, N = 110) = 1.17, *p* > 0.50. Across all three seasons, we found no statistical difference in plant species diversity between environments (Species: U =739, n_1_ = 114, n_2_ =10, p > 0.10, Mean ± SE: Grassland Remnants = 0.77 ± 0.05, Garden = 1.07 ± 0.21). Within the restored native grassland patch, Shannon diversity of biotically-pollinated plant species ranged from 0.25 – 1.96 throughout the entire sampling season (Fig. 3).

**Figure 3.**
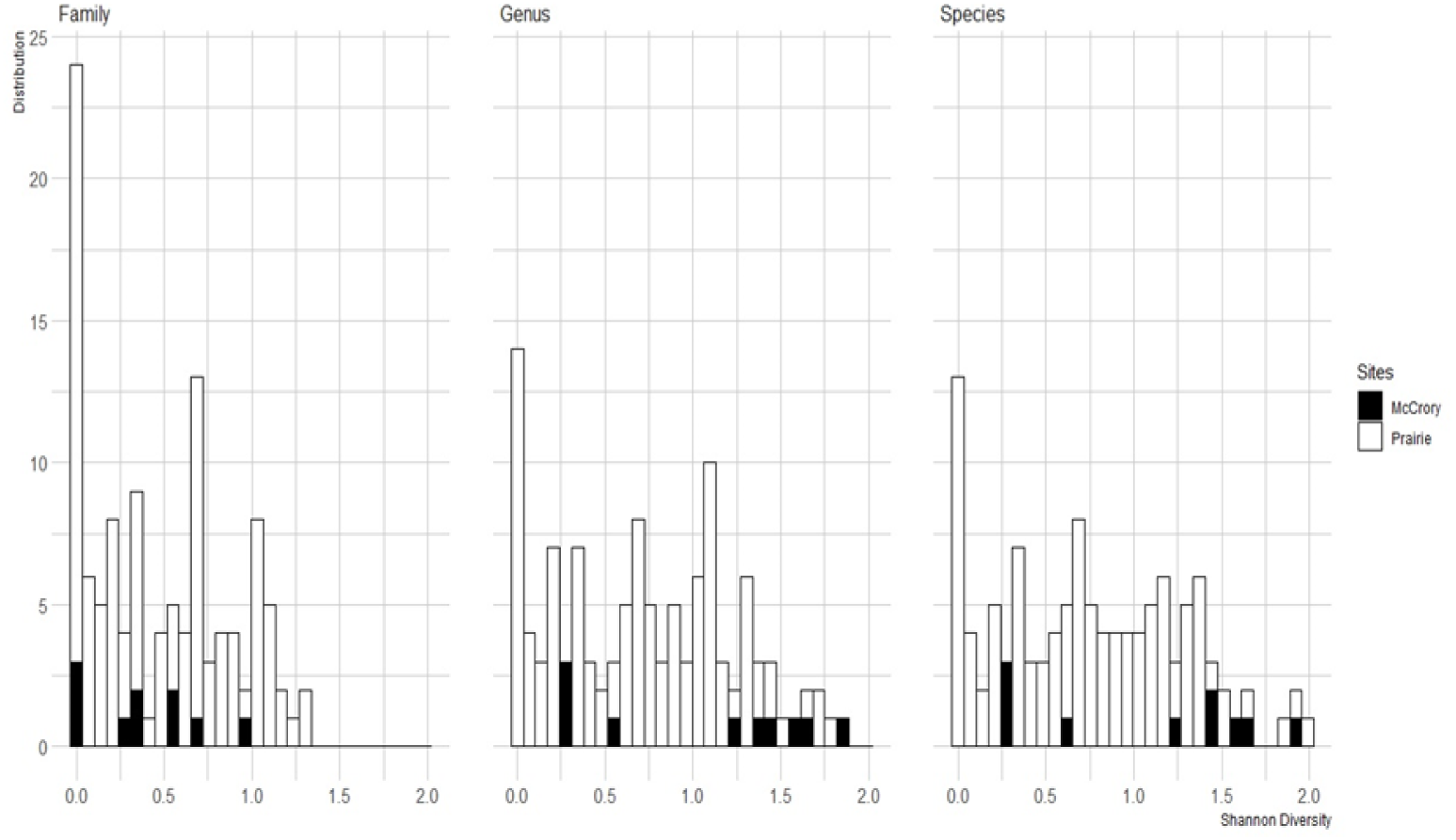
Distribution of Shannon diversity of biotically-pollinated plants at the familial, genera and species level between remnant temperate grassland sites in the Prairie Coteau near Brookings, SD and McCrory Gardens across the three sampling seasons (May – October 2019). The black bars refer to transects conducted in McCrory Gardens while the white bars refer to transects conducted in the remnant temperate grassland sites. Distributions demonstrate diversity on the transect level and are overlayed (i.e., not stacked) for comparison between sites. Height of distribution bars refer to number of samples (transects) that fell within diversity range indicated on the x-axis. Diversity displayed for remnant temperate grassland sites are also provided in Vilella-Arnizaut et al. (bioRxiv, doi.org/10.1101/2021.02.12.431025).

Likewise, Shannon diversity of biotically-pollinated plant species within remnant temperate grassland sites ranged from 0 −2 throughout the entire sampling season. Early season plant species diversity in the restored native grassland patch was 0.25 with *Achillea millefolium* as the most common species recorded on transects (Fig. 4). Within remnant temperate grassland sites, early season plant species diversity ranged from 0 – 1.33 with *Anemone canadensis*, *Gallium boreali* and *Fragaria virginiana* as the most common species found on the transects. Mid-season plant species diversity within the restored native grassland patch ranged from 0.27 - 1.96 with *Coreopsis tictoria* and *Achillea millefolium* as the most common species recorded on transects (Fig. 4). Mid-season plant species diversity within remnant temperate grassland sites ranged from 0 – 2 with *Melilotus sp., Anemone canadensis,* and *Amorpha canescens* as the most common species found on the transects (Fig. 4). Late season plant species diversity within the restored native grassland patch was 1.2 with *Helianthus maximilianii* as the most common species recorded on transects, while late season plant species diversity within remnant temperate grassland sites ranged from 0.17 – 1.5 with *Symphyotrichum lanceolatum*, *Symphyotrichum ericoides* and *Heliopsis helianthoides* as the most common species found on the transects (Fig. 4).

**Figure 4.**
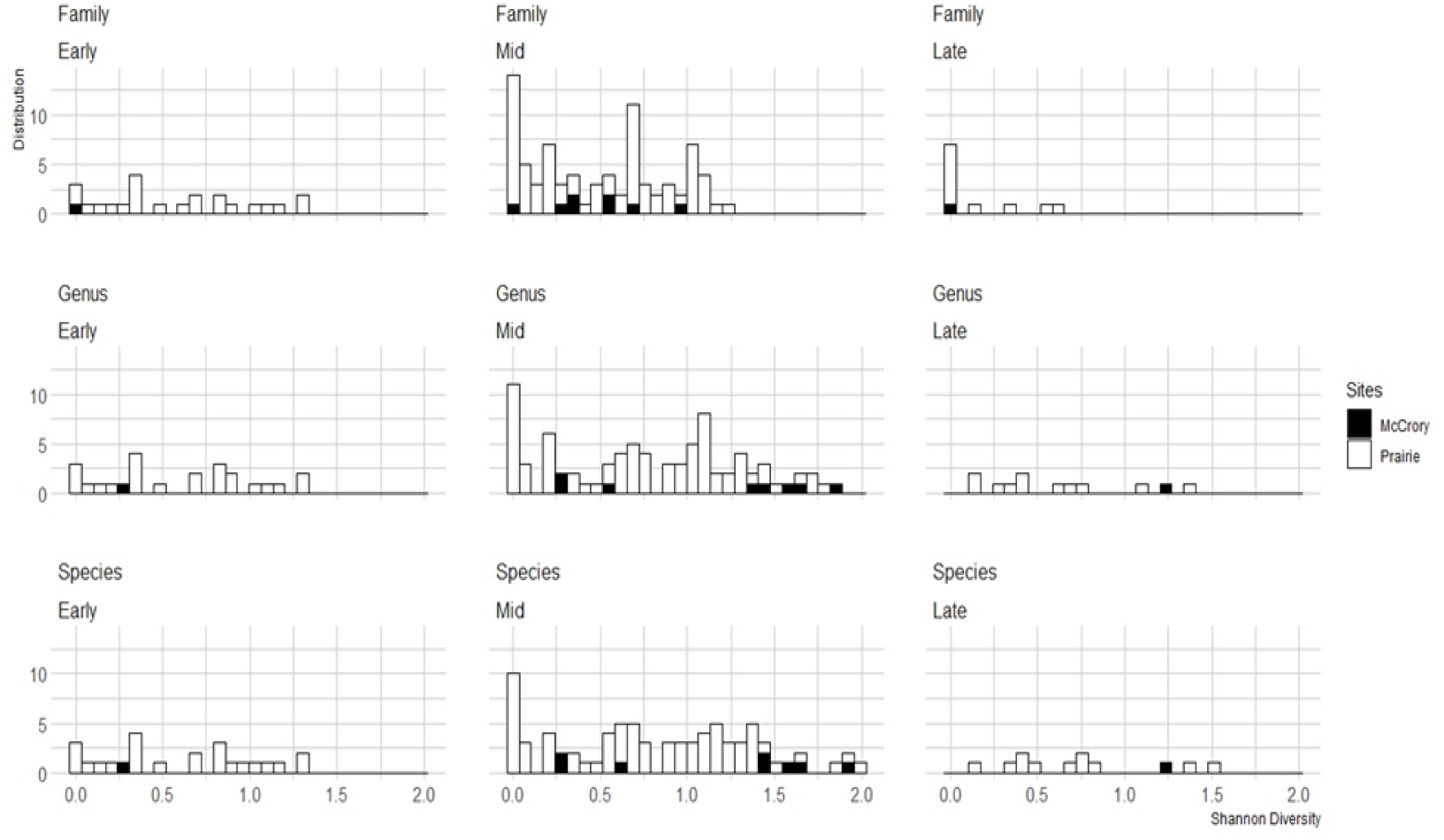
Distribution of Shannon diversity of biotically-pollinated plants at the familial, genera, and species level between remnant temperate grassland sites in the Prairie Coteau near Brookings, SD and McCrory Gardens for each sampling season in 2019 (Early: May – June, Mid: July – August, Late: September – October). The black bars refer to transects conducted in McCrory Gardens while the white bars refer to transects conducted in the remnant temperate grassland sites. Distributions demonstrate diversity on the transect level and are overlayed (i.e., not stacked) for comparison between sites. Height of distribution bars refer to number of samples (transects) that fell within diversity range indicated on the x-axis. Diversity displayed for remnant temperate grassland sites are also provided in Vilella-Arnizaut et al. (bioRxiv, doi.org/10.1101/2021.02.12.431025).

### Plant-pollinator network analysis

Within the restored native grassland patch in McCrory Gardens, we observed 165 unique plant-pollinator interactions and a total of 3,146 observations of pollinators visiting plants from May through October. The most common floral visitors throughout the entire sampling period in McCrory Gardens were Syrphidae (38%), Cantharidae (12%), and Apidae (11%). The plant species with the most interactions in McCrory Gardens throughout the sampling season include *Achillea millefolium* (50%), *Helianthus maximilianii* (8%), and *Solidago rigida* (7.7%). Network specialization (H2’) ranged from 0.26 - 0.64 throughout the entire sampling season for the restored native grassland patch, while network specialization ranged from 0.56 – 0.80 in the remnant temperate grassland communities (Supplemental table 3). Full details for network analyses in the remnant temperate grassland sites are provided in Vilella-Arnizaut et al. (bioRxiv, doi.org/10.1101/2021.02.12.431025), however we provide a synopsis of results below.

Within the restored native grassland patch, connectance ranged from 0.29 – 0.67 and nestedness ranged from 17 – 29 across all three seasons (Supplemental table 3). Likewise, within the remnant temperate grassland communities, connectance ranged from 0.40 – 0.50 and nestedness ranged from 25 – 34 across all three seasons (Supplemental table 3). We did not find a significant difference in H2’ between environments when using transects as our replicates (U = 423, n_1_ =92, n_2_ =10, p > 0.60, Mean ± SE: Grassland Remnants = 0.60 ± 0.03, Garden = 0.57 ± 0.06). Additionally, we found no significant difference in nestedness between environments (U = 495, n_1_ = 92, n_2_ = 10, p > 0.60, Mean ± SE: Grassland Remnants = 26.4 ± 1.72, Garden = 28 ± 1.73), however, we did find the remnant temperate grassland sites to have significantly higher connectance than the restored native grassland patch (U = 250, n_1_ = 92, n_2_ = 10, p < 0.03, Mean ± SE: Grassland Remnants = 0.44 ± 0.016, Garden = 0.34 ± 0.04).

## Discussion

Our study expands on the limited literature available exploring the extent to which botanical gardens can support pollinator communities and pollination services. Previous research has examined how urbanization and impervious surfaces may impact pollinator movement (Fortel et al. 2014; Levé et al. 2019). Recent work has highlighted the potential conservation value of urban green spaces for pollinator communities, especially those found within cities (Micholap et al. 2017; Lewis et al. 2019). We further develop these approaches by quantifying and comparing the diversity and interactions of plant-pollinator communities within a restored native grassland patch centered in a botanical garden and remnant temperate grassland habitats in order to understand how these environments may differ with regards to plant-pollinator community structure. We found that the restored native grassland patch in McCrory Gardens fell within similar ranges of Shannon diversity for both plant and pollinator communities in comparison to those found within remnant temperate grasslands. Network metrics were similar across seasons between communities, except for connectance. Below, we discuss and compare the diversity and network community structure between remnant temperate grassland habitats and the restored native grassland patch in McCrory Gardens.

### Comparing *and* contrasting diversity of the plant-pollinator communities

Across all three seasons, pollinator diversity in all taxonomic groups within the restored native grassland patch overlapped with the mid to upper range of diversity reflected in the remnant temperate grassland transect samples (Fig. 1). Pollinator diversity within the restored native grassland patch was greater than 55% of total remnant temperate grassland transects across the three sampling seasons. For both remnant temperate grassland communities and the restored native grassland patch, pollinator diversity was greatest in the mid and late seasons. These results for pollinator community diversity indicate the restored native grassland patch in the botanical garden can maintain a relatively diverse pollinator community comparable to the diversity found within remnant temperate grassland habitats in the same region. Greater pollinator diversity from genus to functional group level could benefit botanical gardens and urban green spaces by promoting community resiliency through functional redundancy (Kühsel and Blüthgen 2015).

Likewise, floral community diversity within the restored native grassland patch overlapped with the mid to upper range of diversity reflected in the remnant temperate grassland transect samples across all three seasons. However, we noted the restored native grassland patch is less diverse than remnant temperate grasslands in the early season. Approximately 90% of the transects sampled in remnant temperate grassland sites had greater floral diversity in all taxonomic groups (e.g., family, genus, species) in the early season compared to the restored native grassland patch. However, when compared across all three seasons, we did not find a significant difference between environments. The increased floral genus and species diversity found within the restored native grassland patch in the mid and late season (Fig. 4) is likely due to the diversity within Asteraceae, as approximately 96% of the individuals we documented in the garden transects belong to this family. This may also explain why floral diversity in the restored native grassland patch is lower across all taxonomic groups in the early season, as the vast majority of asters we sampled bloomed in the mid and late seasons.

### Comparing and contrasting network metrics

The greatest overlap in network metrics (i.e., nestedness, connectance, and specialization) between the restored native grassland patch and remnant temperate grasslands occurred during the mid-season. Across seasons, indices for nestedness and network specialization demonstrated no significant difference. However, values for connectance were significantly higher in the remnant temperate grassland sites than the restored native grassland patch. Connectance is often used in ecological networks to measure community complexity and is generally positively associated with conservation value (Dunne et al. 2002; Thébault and Fontaine 2010; Tylianakis et al. 2010; Hagen et al. 2012). Communities with increased interaction complexity are expected to be more stable and robust to species loss in theory (Dunne et al. 2002). However, Heleno (et al. 2012) notes that connectance alone should not be used to determine conservation value as it is context-specific depending on the different conservation values of species in a network. Overall, results indicate that plant-pollinator community interactions in the restored native grassland patch are less complex than remnant temperate grassland sites. The higher level of complexity in plant-pollinator communities within natural habitats may be attributed to the distinct phenological shifts in the flowering community across seasons, which have evolved with the local pollinator fauna over a longer evolutionary time scale (Gomez & Zamora 2006; Minckley & Roulston 2006; Craine et al. 2012). This temporal variability could explain how natural habitats maintain more complex interactions than their garden counterparts. Successful recruitment of native plants is an on-going challenge in temperate grasslands (Martin and Wilsey 2006; Gibson-Roy et al. 2007; Johnson et al. 2018) and may be an obstacle botanical gardens will have to overcome when seeking to maintain complex and stable plant-pollinator communities in restoration plots. Botanical gardens that wish to establish native plant restoration plots will likely need to consider a balance between aesthetics and diversity in order to increase the complexity of plant-pollinator community interactions.

Moreover, the landscape surrounding natural habitats may provide other resources (e.g., nesting resources) that some pollinators may require in order to thrive, particularly those whose foraging distance is shorter to other more generalized and mobile visitors (i.e., honey bees) (Beekman and Ratnieks 2000). The spatial variability of resources found within natural habitats is likely a factor attributing to the difference in connectance between environments, though landscape analysis for the garden community was beyond the scope of this paper. In general, the restored native grassland patch within McCrory Gardens demonstrates similar plant-pollinator community structure to the remnant temperate grassland sites. Nested networks displaying a higher degree of connectance are considered more resilient and stable, making them important considerations for conservation value (Memmott et al. 2004; Okuyama and Holland 2008; Thébault and Fontaine 2010). The nested pattern found in the networks indicates a degree of interaction redundancy that likely contributes to community stability (Bascompte et al. 2003; Nielsen and Bascompte 2007). However, it appears that the remnant temperate grassland habitats within the Northern Great Plains support a greater degree of interaction complexity in their plant-pollinator communities. This could be concerning for maintaining stable pollination services in botanical gardens, as community complexity is associated with stable and robust communities.

## Conservation Implications

Our findings demonstrate the promising role botanical gardens could play as restoration reservoirs for local pollinator communities by supporting plant-pollinator interactions comparable to those found in natural habitat remnants in the same region. In the absence of large swaths of preserved habitat, small reservoirs have been notably valuable for wildlife conservation, though the context of the landscape is important when seeking to maximize regional insect diversity (Shafer 1995; Tscharntke et al. 2002). Though this study does not directly examine landscape effects which may explain some differences between environments, the restored native grassland patch located in McCrory Gardens demonstrated comparable measures of plant-pollinator community structure to natural habitats and greater pollinator diversity indicating the garden’s potential in serving as a beneficial patch for pollinator communities. Future work studying the influence of increased green spaces in urban areas in conjunction with conserving remaining patches of natural habitat will be invaluable in our understanding of how best to conserve pollinator communities and stable pollination services.

Temperate grasslands are among the least protected habitat types in the world, with conversion outpacing conservation by eight to one (Hoekstra et al. 2005). In the United States, the temperate grasslands of the Northern Great Plains are a valuable resource for approximately 40% of transported honey bee colonies from May through October by providing abundant floral resources through regional blooms (USDA, 2014). However, the Great Plains has experienced considerable habitat loss due to landscape conversion with more than 96% of the grassland habitat of the Great Plains already been converted to cropland or other less diverse vegetation (Bauman et al. 2016). Botanical gardens have the potential to provide abundant floral resources to pollinator communities within increasingly disturbed landscapes; however, the role of botanical gardens in pollinator conservation is critically understudied.

Our aim for this study was to provide further understanding on the extent to which botanical gardens can serve as supplementary resources for pollinator communities within critically fragmented landscapes. More research focused on plant-pollinator interactions in botanical gardens, particularly in regions that experience distinct flowering shifts within the growing season, paired with sampling of plant-pollinator interactions in natural habitats could help us understand how effective botanical gardens may be as additional sources of habitat. Increasing sampling within distinct flowering seasons and environments could provide important context for conservation of pollination services on a wider scale. For example, within the restored native grassland patch in McCrory gardens, we found that floral diversity is similar to floral diversity in the remnant temperate grasslands, however, we see that floral diversity within the restored native grassland patch is primarily from Asteraceae. Extending sampling to include early season species could elucidate how early season pollinators may be affected by this gap in resources before Asteraceae species are blooming. Consequently, gardens could adjust management once these nuances are better understood. Additionally, extending research across multiple years could provide valuable insight into how plant-pollinator communities may shift following the progression of native restoration gardens. Continued research tracking the influence and progression of green spaces on plant-pollinator interactions over time could expand as initiative for private and public green spaces grows. Increasing urban garden areas may very well act similarly to habitat corridors, which have been shown to be beneficial in improving wildlife conservation (Correa Ayram et al. 2016). By understanding the effectiveness of botanical gardens in supporting pollinator populations, we can view urban spaces as valuable conservation tools rather than barriers.

## ACKNOWLEDGMENTS

South Dakota State University is located on the ancestral territory of the Oceti Sakowin [oh-CHEH-tee SHAW-koh-we], the proper name for the people commonly called Sioux. We thank S. Stiles, J. Gelderman, T. Wallner, N. Petersen, E. Baier, and S. Daniels for field assistance, J. Purintun for providing botanical expertise, the staff at McCrory Gardens for their support, and the landowners and managers that provided permission and advice on site selection, including South Dakota Game Fish and Parks, U.S. Fish and Wildlife Services, The Nature Conservancy, and the City of Brookings. This project was funded from several Hatch grants and the North Central Sun Grant Initiative (USDA/DOE) SA1500640.

## Appendices

## Appendix A

The table below lists all of the pollinators observed and identified in the Prairie Centennial Garden within McCrory Gardens, Brookings, South Dakota in 2019 down to lowest taxonomic level. Insect pollinators that could not be identified to genus were placed in a catch-all genus that consisted of the first five letters of their family name. For example, for a fly pollinator in the family Muscidae, we created a genus named Gen_Musci in the dataset in order to include these visitors in the analyses. All insect pollinators could be identified down to family except for two non-syrphid fly visitors we named Gen_Myst which make up < 0.5 % of visitations. Table also includes number of total observed interactions of each pollinator from May through October 2019.

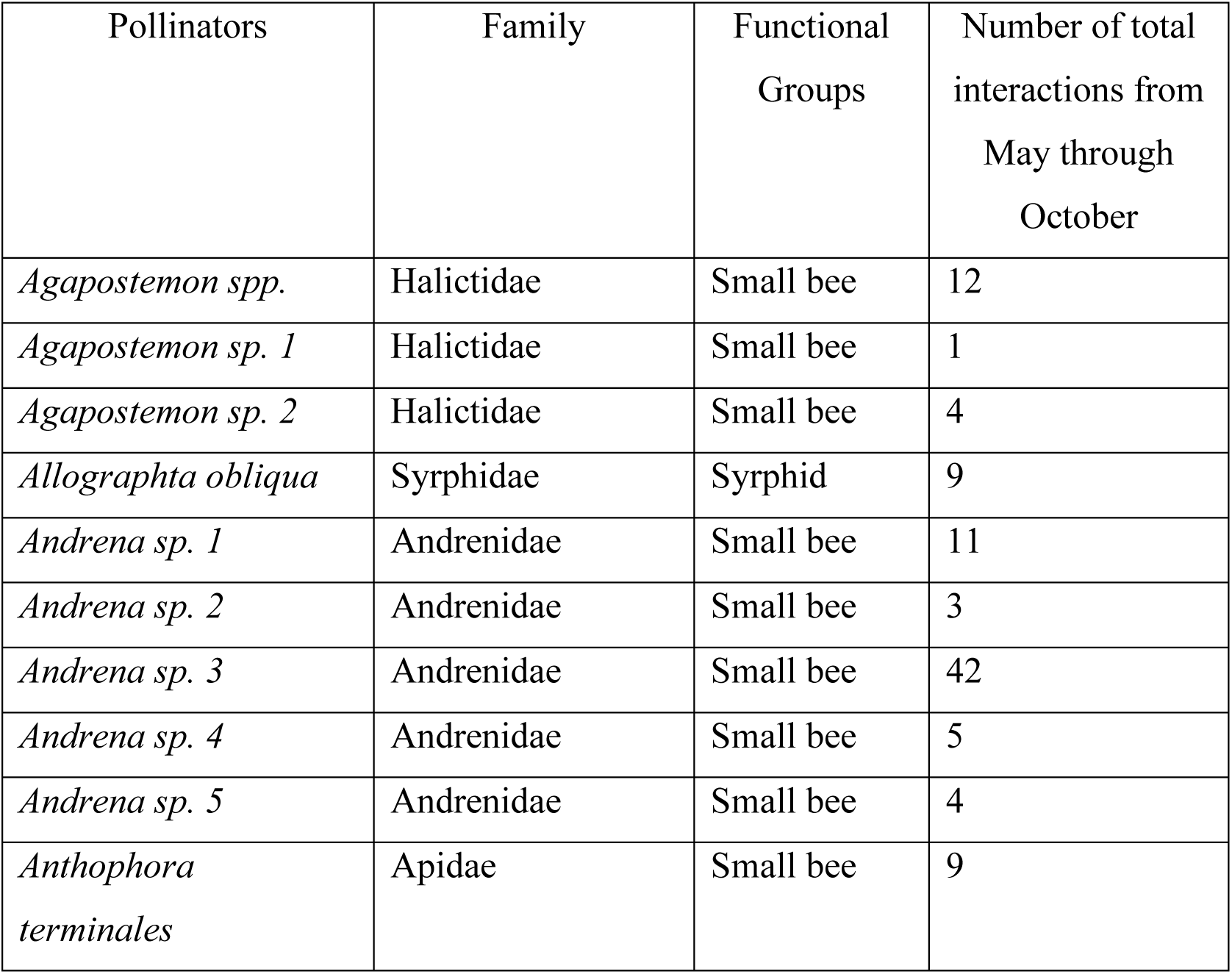
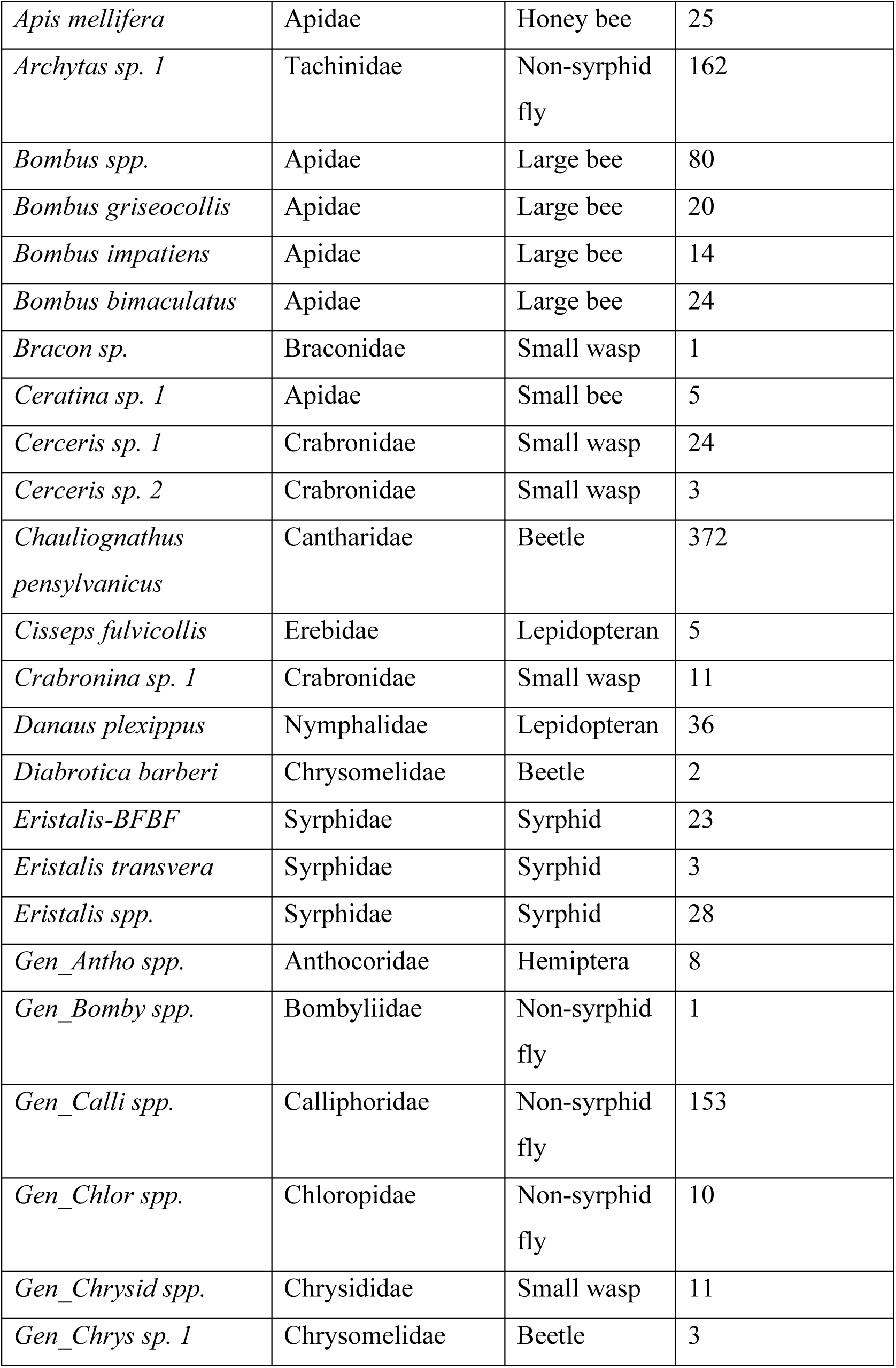
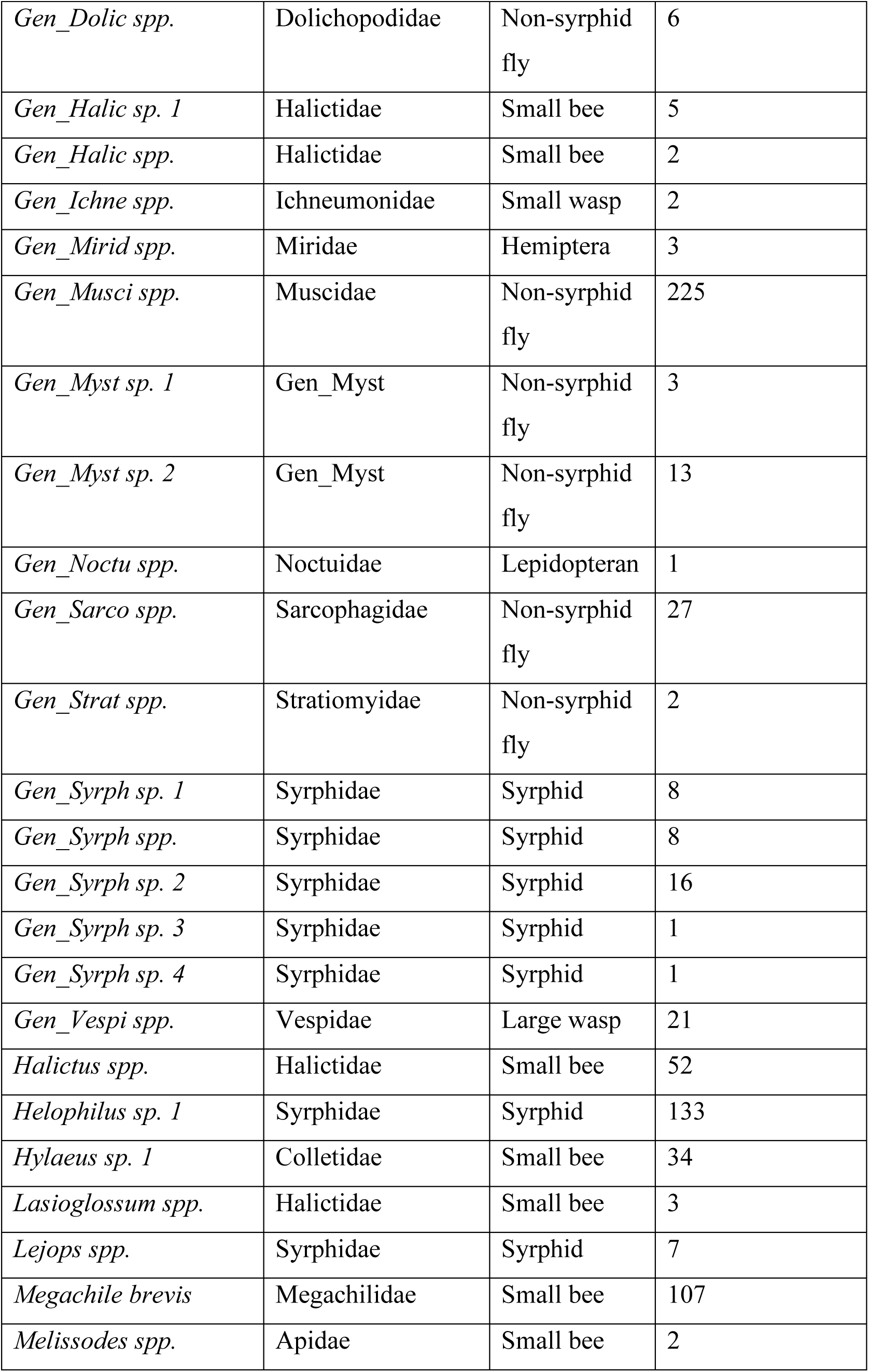
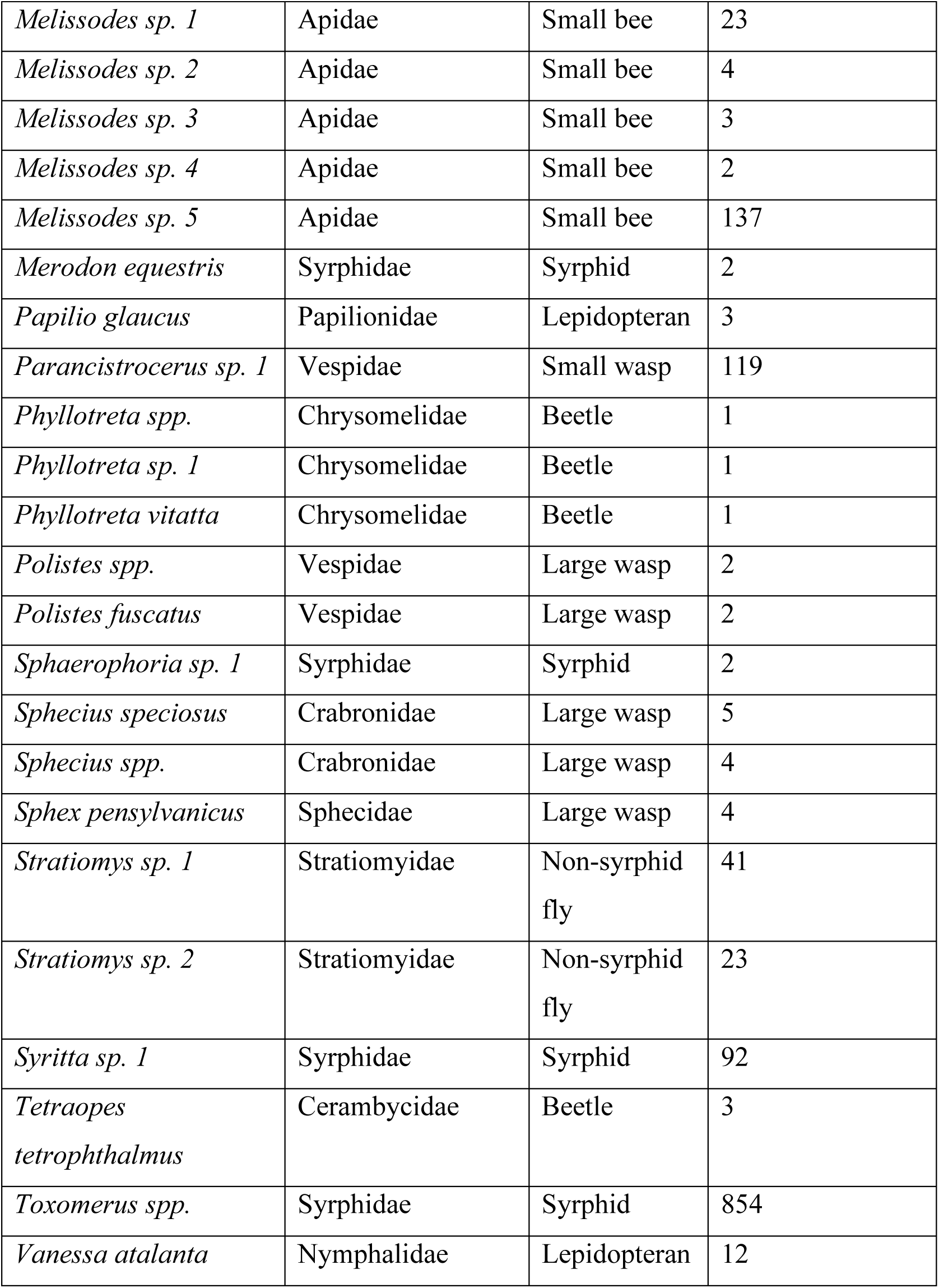

## Appendix B

The table below lists all of the biotically-pollinated flowering plants identified in the Prairie Centennial Garden within McCrory Gardens, Brookings, South Dakota in 2019 down to lowest taxonomic level. All species except Epilobium sp. were identified to species level.

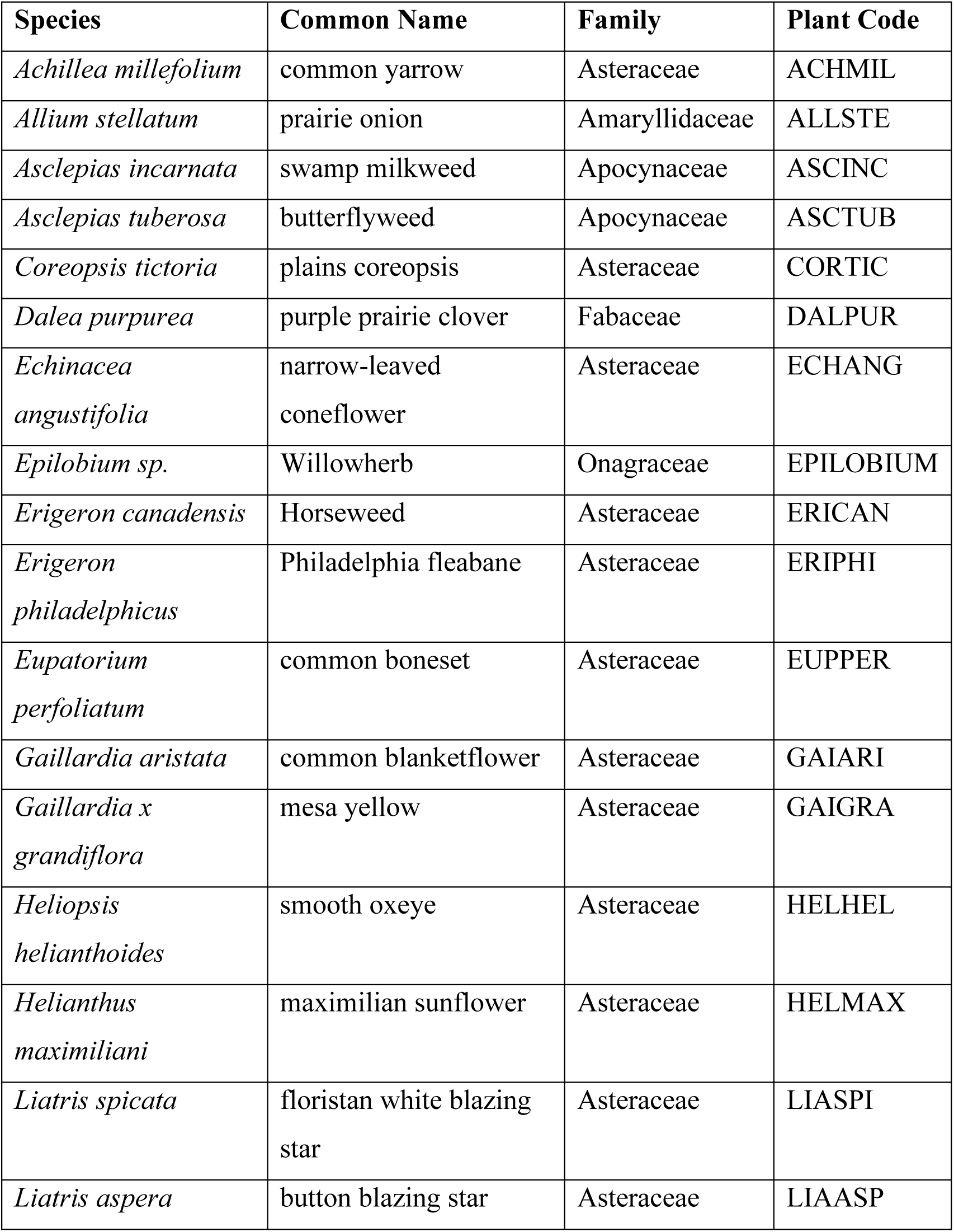
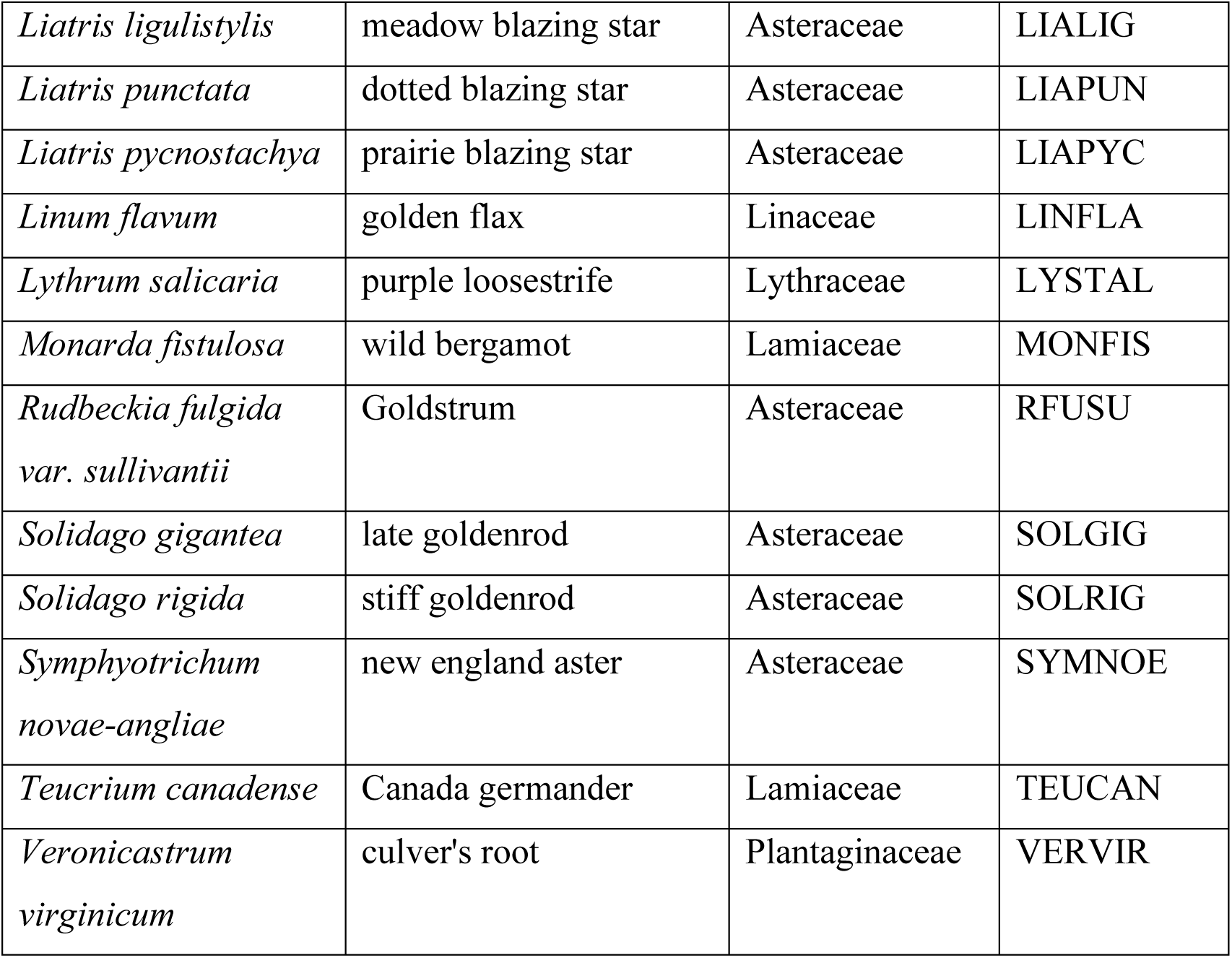

## Notes

### Competing Interest Statement

The authors have declared no competing interest.

