## Supplementary Materials Figures and Tables for "Use of botanical gardens as arks for conserving pollinators and plant-pollinator interactions: A case study from the US Northern Great Plains"

Table 1. Table of Spearman rank correlations for pollinator and plant diversities with Rho values and Bonferroni-corrected p values from early (May-June), mid (July-August), and late (September-October) seasons in the Prairie Centennial Garden in McCrory Gardens, Brookings, South Dakota in 2019. *P* < 0.05; * *P* < 0.05; ** *P* < 0.01; *** *P* < 0.001

| **Season** | **Metrics tested** | **p-value** | **Rho value** |
| --- | --- | --- | --- |
| **Total** | Pollinator Functional Diversity & Pollinator Family Diversity | 0.17 | 0.48 |
| **Total** | Pollinator Functional Diversity & Pollinator Genus Diversity | 0.19 | 0.45 |
| **Total** | Pollinator Family Diversity & Pollinator Genus Diversity | 0.28 | 0.38 |
| **Early** | Pollinator Functional Diversity & Pollinator Family Diversity | 0.17 | 0.48 |
| **Early** | Pollinator Functional Diversity & Pollinator Genus Diversity | 0.19 | 0.45 |
| **Early** | Pollinator Family Diversity & Pollinator Genus Diversity | 0.28 | 0.38 |
| **Mid** | Pollinator Functional Diversity & Pollinator Family Diversity | 0.17 | 0.48 |
| **Mid** | Pollinator Functional Diversity & Pollinator Genus Diversity | 0.19 | 0.45 |
| **Mid** | Pollinator Family Diversity & Pollinator Genus Diversity | 0.28 | 0.38 |
| **Late** | Pollinator Functional Diversity & Pollinator Family Diversity | 0.17 | 0.48 |
| **Late** | Pollinator Functional Diversity & Pollinator Genus Diversity | 0.19 | 0.45 |
| **Late** | Pollinator Family Diversity & Pollinator Genus Diversity | 0.28 | 0.38 |
| **Total** | Floral Family Diversity & Floral Genus Diversity | 0.001 | 0.87 |
| **Total** | Floral Family Diversity & Floral Species Diversity | 0.001 | 0.87 |
| **Total** | Floral Genus Diversity & Floral Species Diversity | 2.20E-16 | 1 |
| **Early** | Floral Family Diversity & Floral Genus Diversity | 0.001 | 0.87 |
| **Early** | Floral Family Diversity & Floral Species Diversity | 0.001 | 0.87 |
| **Early** | Floral Genus Diversity & Floral Species Diversity | 2.20E-16 | 1 |
| **Mid** | Floral Family Diversity & Floral Genus Diversity | 0.001 | 0.87 |
| **Mid** | Floral Family Diversity & Floral Species Diversity | 0.001 | 0.87 |
| **Mid** | Floral Genus Diversity & Floral Species Diversity | 2.20E-16 | 1 |
| **Late** | Floral Family Diversity & Floral Genus Diversity | 0.001 | 0.87 |
| **Late** | Floral Family Diversity & Floral Species Diversity | 0.001 | 0.87 |
| **Late** | Floral Genus Diversity & Floral Species Diversity | 2.20E-16 | 1 |

Table 2. Mann-Whitney U test results comparing pollinator functional group diversity, pollinator family diversity, floral family diversity, and floral genus diversity across all seasons using transects as our replicates between McCrory Gardens and prairies sites in the Prairie Coteau region near Brookings, SD in 2019.

| **Diversity** | **U statistic** | **N_1_** | **N_2_** | **P value** | **Mean ± SE of Prairies** | **Mean ± SE of Garden** |
| --- | --- | --- | --- | --- | --- | --- |
| Pollinator Functional Groups | 925 | 114 | 10 | 0.001 | 0.86 ± 0.04 | 1.23 ± 0.07 |
| Pollinator Family | 927 | 114 | 10 | 0.001 | 1.014 ± 0.04 | 1.48 ± 0.08 |
| Floral Family | 491 | 114 | 10 | 0.47 | 0.48 ± 0.04 | 0.37 ± 0.10 |
| Floral Genus | 763 | 114 | 10 | 0.08 | 0.71 ± 0.05 | 1.05 ± 0.20 |

Table 3. Means of network metrics across all prairie sites in the Prairie Coteau region near Brookings, SD and network metrics of McCrory Gardens for all seasons using transects as our replicates in 2019 (Early: May – June, Mid: July – August, Late: September - October). Standard errors of means are placed in parentheses next to mean value. McCrory Gardens was only sampled once during the early and late seasons; thus we do not provide standard errors for these values.

| **Site** | **Nestedness** | **Connectance** | **Network Specialization** | **Season** |
| --- | --- | --- | --- | --- |
| Prairies | 25 (4.5) | 0.50 (0.03) | 0.56 (0.07) | Early |
| McCrory | 28 | 0.38 | 0.26 | Early |
| Prairies | 26 (1.9) | 0.40 (0.02) | 0.60 (0.04) | Mid |
| McCrory | 29 (1.6) | 0.29 (0.02) | 0.64 (0.05) | Mid |
| Prairies | 34 (7.4) | 0.45 (0.05) | 0.80 (0.13) | Late |
| McCrory | 17 | 0.67 | 0.35 | Late |
